# Biochemical characterization of cyanobacterial HtpG from *Synechococcus elongatus* PCC 7942

**DOI:** 10.1101/2025.04.03.647111

**Authors:** Liqun Jiang, Ibrahim D. Boyenle, Nicolas Delaeter, Yanxin Liu

**Affiliations:** Institute for Bioscience and Biotechnology Research, University of Maryland, Rockville, MD 20850, USA; Department of Chemistry and Biochemistry, University of Maryland, College Park, MD 20740, USA

**Keywords:** cyanobacterium, HtpG, Hsp90, chaperone, ATPase, protein homeostasis, stress response

## Abstract

The 90 kDa Heat Shock Protein (Hsp90) is an essential and highly conserved molecular chaperone that supports the folding and maturation of a diverse array of client proteins across prokaryotic and eukaryotic organisms. In bacteria, HtpG, the Hsp90 homolog, plays a central role in stress response and protein homeostasis, particularly under high-temperature and other stress conditions. Despite extensive studies on HtpG from *E. coli*, the biochemical properties and functional roles of cyanobacterial HtpG remain poorly characterized. Here, we focus on HtpG from the cyanobacterium *Synechococcus elongatus* PCC 7942 (seHtpG), a model organism for photosynthesis and circadian rhythm research. We developed a method for the overexpression and purification of seHtpG in *E. coli*, achieving high purity and yield suitable for biochemical and structural studies. Biophysical and biochemical assays show that seHtpG forms dimers and hydrolyzes ATP and a rate of 1.8 ATP/min, faster than that of *E. coli* HtpG. This work establishes seHtpG as a model for studying the roles of HtpG in cyanobacterial protein homeostasis, photosynthesis, and stress response, enabling further exploration of cyanobacterial Hsp90 in ecosystem dynamics and biotechnological applications.

**Highlights:**

1. HtpG from *S. elongatus* PCC 7942 (seHtpG) was recombinantly overexpressed.
2. seHtpG was purified to high homogeneity using chromatography techniques.
3. Like other members of the Hsp90 family, seHtpG forms a dimeric structure.
4. seHtpG exhibits ATPase activity of 1.8 ATP/min at 37°C.

## 1. Introduction

The 90 kDa Heat Shock Protein (Hsp90) is a highly conserved and ubiquitous molecular chaperone essential for the folding and maturation of a wide range of client proteins in both prokaryotic and eukaryotic cells [1, 2]. In humans, Hsp90 stabilizes many oncogenic proteins—including kinases, hormone receptors, and transcription factors—that are crucial for signal transduction and cellular regulation [3]. In bacteria, HtpG (high-temperature protein G) serves as the Hsp90 homolog and is central to bacterial stress responses and protein homeostasis, especially under high-temperature or other stress conditions [4, 5]. Both *E. coli* HtpG and yeast Hsp90s have been extensively studied as model systems for the Hsp90 family, while the therapeutic potential of human Hsp90s has driven significant research interest [1, 6]. In addition to its presence in *E. coli*, the gene encoding HtpG has been identified in cyanobacteria [7, 8]. However, the biochemical properties of cyanobacterial HtpG, and its roles in photosynthesis and stress response, remain poorly understood.

Cyanobacteria are prokaryotic, oxygenic phototrophs with broad ecological distribution, found in nearly every conceivable habitat on Earth, including extreme environments such as bare rocks and highly toxic wastewater [9]. Cyanobacteria drive ecosystem nutrient cycling through photosynthesis, fundamentally contributing to global carbon and nitrogen fixation [10]. Beyond their ecological roles, cyanobacteria are rich sources of bioactive compounds, making them valuable in bioremediation and as sources of biofertilizers, biofuels, food additives, and pigments [11, 12]. Cellular protein homeostasis in cyanobacteria is crucial for ecosystem stability, especially under environmental stress, and relies on a complex chaperone network, with HtpG playing a major role in this system [7, 8, 13, 14].

In this study, we investigate the cyanobacterial HtpG from *Synechococcus elongatus* PCC 7942, a model organism widely used for research on prokaryotic photosynthesis, circadian rhythms, and as a biotechnological platform [15, 16]. The genome of *Synechococcus elongatus* PCC 7942 contains a single gene encoding HtpG (Uniprot ID Q79N42; referred to hereafter as seHtpG). Sequence alignments between seHtpG and other Hsp90 family members reveal high sequence and structural conservation. seHtpG comprises 638 amino acids and contains a conserved ATP-binding pocket in its N-terminal domain (NTD), supporting its predicted ATPase activity. A conserved dimerization interface in the C-terminal domain (CTD) suggests that seHtpG forms a dimer, while the middle domain (MD), connecting the NTD and CTD, is also conserved and can be further divided into Large-MD and Small-MD regions.

To investigate the structure and function of seHtpG, one has to recombinantly overexpress and purify seHtpG to homogeneity. Here, we report our successful production and purification of recombinant seHtpG in *E. coli* using high-performance liquid chromatography. We further characterize its oligomerization state and ATPase activity, laying a foundation for future studies on the role of HtpG in protein homeostasis, photosynthesis, and stress response in cyanobacteria.

## 2. Materials and Methods

### 2.1 Sequence analysis

The sequence alignment to show the identity and similarity between Hsp90 homologs were analyzed using the Clustal W program (http://www.ebi.ac.uk/Tools/msa/clustalw2/). The Hsp90 sequences were obtained from Uniprot database and include the following Hsp90s homologs with abbreviation and Uniprot ID shown in the parentheses: HtpG from *Synechococcus elongatus* PCC 7942 (seHtpG, Q79N42), HtpG from *Synechococcus* sp. PCC 7002 (ssHtpG, B1XQJ4), HtpG from *E. coli* (ecHtpG, P0A6Z3), Hsc82 from *Saccharomyces cerevisiae* (scHsc82, P15108), ⍰- and β-isoforms of cytosolic Hsp90 from human (hsHsp90⍰, P07900; hsHsp90β, P08238), mitochondrial Hsp90 from human (hsTrap1, Q12931), endoplasmic reticulum Hsp90 from human (hsGrp94, P14625), cytosolic Hsp90 from *Arabidopsis thaliana* (atHsp901, P27323), chloroplastic Hsp90 from *Arabidopsis thaliana (*atHsp905, Q9SIF2), and Hsp90 form *Chlamydomonas reinhardtii* (crHsp90, A8J1U1). The Phylogenetic Tree was calculated using neighbor-joining method on Mega 11.

### 2.2 Protein Expression

The DNA sequence of seHtpG was codon-optimized for expression in *E. coli* and cloned into a pET28a expression vector, incorporating an N-terminal 6xHis tag followed by a TEV protease cleavage site. Protein expression was carried out in *E. coli* BL21(DE3)-RIL and Rosetta (DE3) cells, which were grown at 37°C with shaking at 150 rpm to an OD600 of 0.4–0.6. For induction, the temperature was maintained at 37°C or reduced to 16°C, followed by the addition of 0.5 mM IPTG. Protein expression proceeded for 3 hours at 37°C or overnight at 16°C.

### 2.3 Protein purification

*E. coli* cells were harvested by centrifugation and resuspended in buffer containing 40 mM Tris·HCl (pH 7.5), 480 mM KCl, 20 mM imidazole, and 1 mM β-mercaptoethanol. Lysis was performed by homogenization and sonication, followed by centrifugation and filtration through a 0.45 μm membrane.

Initial purification of seHtpG was conducted using Ni-NTA affinity chromatography. The elution from the Ni-NTA column was dialyzed overnight to remove imidazole, with TEV protease added to cleave the 6xHis tag. Further purification was achieved using ion exchange chromatography (MonoQ 10/100 GL column, GE Life Sciences) and size exclusion chromatography (SEC) (Superdex 200 16/60 column, GE Life Sciences). Elution profiles were monitored as absorbance at 280 nm, and the efficiency of each purification step was assessed via 4-12% SDS-PAGE. Protein concentrations were determined spectroscopically using the molar extinction coefficient (ε), calculated based on the amino acid sequence.

### 2.4 Analytical ultracentrifugation

Analytical ultracentrifugation (AUC) was conducted using a Beckman Optima XL-A analytical ultracentrifuge. Prior to AUC runs, seHtpG was dialyzed overnight at 4°C in buffer containing 40 mM HEPES–HCl (pH 7.5), 150 mM KCl, and 1 mM β-mercaptoethanol. AUC experiments were performed at 45,000 rpm (AN-60Ti rotor) for 8–10 hours at 20°C, with absorbance monitored at 280 nm. Scans were collected at approximately 60-second intervals with a radial step size of 0.001 cm. Sedimentation velocity analysis was carried out using SEDFIT/SEDPHAT (NIH) software to determine the standard sedimentation coefficients (s20,w). Buffer viscosity (η = 1.0254 × 10□^2^ poise), buffer density (ρ = 1.00847 g/mL), and the partial-specific volume of seHtpG (0.7300 mL/g at 20°C, as a dimer) were calculated using the Sednterp program (www.jphilo.mailway.com/download.htm).

### 2.5 Steady-state enzymatic coupled ATPase assay

The steady-state ATP hydrolysis rate of seHtpG was measured using an enzymatic coupled ATPase assay. The final assay mixture contained 40 mM HEPES (pH 7.5), 150 mM KCl, 1 mM MgCl□, 1 mM ATP, 0.6 mM NADH, 1 mM phosphoenol pyruvate, 25 U/mL pyruvate kinase, 25 U/mL lactate dehydrogenase, and 4–20 µM seHtpG monomer. NADH consumption was monitored by measuring absorbance at 340 nm on a Spectramax iD5 plate reader (Molecular Devices, LLC, USA). The absolute ADP release rates were calculated using a standard curve correlating ADP concentration with the decrease in absorbance at 340 nm.

## 3. Results and discussion

### 3.1 Sequence analysis

To explore the evolution of cyanobacterial HtpG, we analyzed the protein sequence of HtpG from *Synechococcus elongatus* PCC 7942 (seHtpG, UniProt ID: Q79N42) in comparison with orthologs from *Synechococcus sp*. PCC 7002, *Chlamydomonas reinhardtii* (microalgae), *Arabidopsis thaliana* (plant), *E. coli* (bacteria), *Saccharomyces cerevisiae* (yeast), and *Homo sapiens* (human). Note that the genome of *S. elongatus* PCC 7942 contains only a single copy of the *htpg* gene (NCBI: NZ_JACJTX010000001.1).

Sequence alignment reveals a high degree of conservation, particularly within cyanobacterial species (Fig. 1A). Like other Hsp90s, seHtpG includes three conserved domains: the N-terminal domain (NTD), middle domain (MD), and C-terminal domain (CTD). However, seHtpG lacks the charged linker between the NTD and MD, as well as the MEEVD motif at the C-terminus, both of which are characteristic of cytosolic Hsp90s in eukaryotes (Fig. 1A). This absence of the charged linker and MEEVD motif is also observed in *E. coli* HtpG and the human mitochondrial Hsp90, Trap1. Notably, cyanobacterial HtpG contains a unique 10-30 residue insertion (E391 to T408 in seHtpG, Fig. 1A) between the large and small MD segments, a feature absents in non-cyanobacterial species. Although the sequence and length of this insertion vary, its presence is conserved across cyanobacteria. The structural and functional implications of this insertion are yet to be determined.

**Figure 1.**
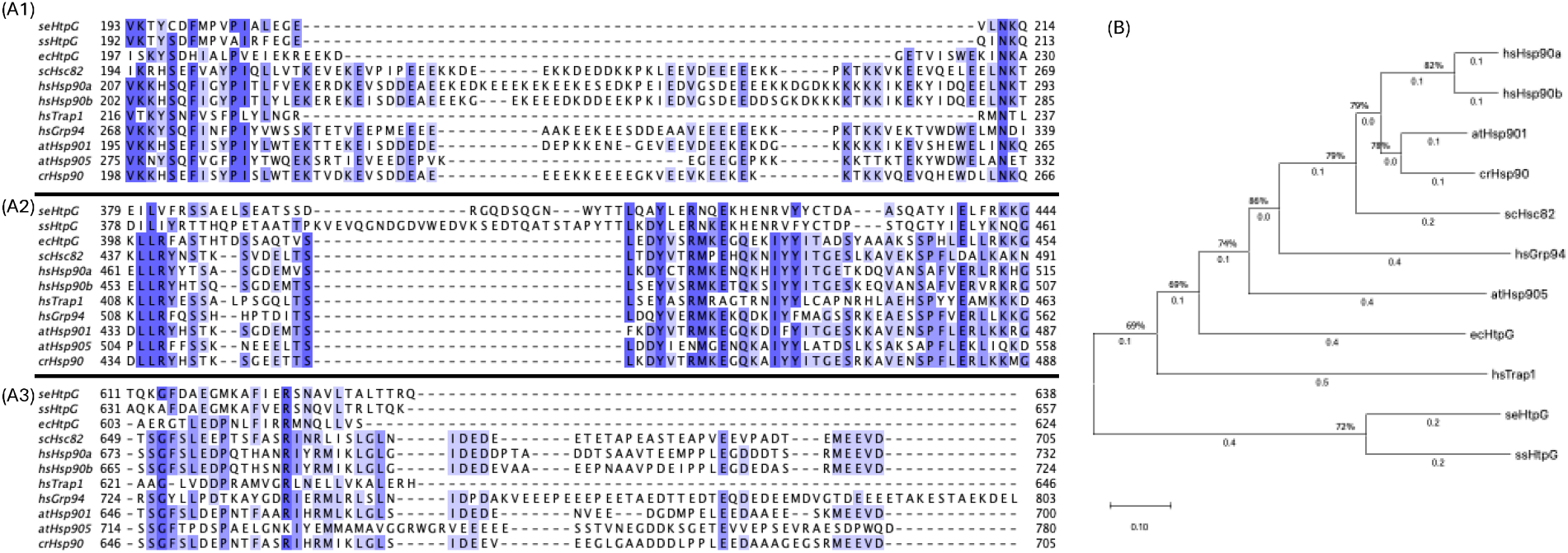
Sequence alignment and phylogenetic analysis of Hsp90 family members across prokaryotic and eukaryotic organisms. (A) Partial sequence alignment highlights key differences between Synechococcus elongatus PCC 7942 HtpG (seHtpG) and other Hsp90 homologs. Conserved regions and unique sequence features in seHtpG are indicated. (B) Phylogenetic tree depicting relationships among representative Hsp90 homologs, illustrating the close evolutionary relationship between cyanobacterial HtpG and mitochondrial Hsp90 (Trap1) from eukaryotes. Numbers on internal branches represent bootstrap support values, indicating the reliability of branching patterns. A branch length of 0.10 represents the phylogenetic distance in terms of sequence divergence.

Photosynthetic cyanobacteria are widely regarded as the evolutionary origin of chloroplasts in eukaryotic plant cells [17]. Consequently, we anticipated that seHtpG would be phylogenetically close to Hsp90 homologs in algae or higher plants, particularly the chloroplastic Hsp90. However, *seHtpG* displays its closest phylogenetic relationship with the human mitochondrial Hsp90, Trap1 (Fig. 1B). Mitochondria, like chloroplasts, are thought to have originated from an ancient endosymbiotic event. These findings suggest a potential common ancestry between the cyanobacterium-like ancestor of chloroplasts and the bacterium-like ancestor of mitochondria, indicating possible shared evolutionary origins between these organelles.

### 3.2 Recombinant overexpression of seHtpG in *E. coli*

To optimize overexpression of recombinant seHtpG in *E. coli*, we tested various expression and induction conditions, including *E. coli* strains, IPTG induction temperature, and induction duration. Results were analyzed by SDS-PAGE with Coomassie staining (Fig. 2). A prominent band between 65 and 80 kDa indicates successful overexpression of seHtpG, which has a theoretical molecular weight of 72.6 kDa for the wild-type protein, or 76 kDa for the fusion protein including the 6xHis tag, TEV cleavage site, and linkers. We evaluated two *E. coli* strains, Rosetta (DE3) and BL21 (DE3)-RIL, for seHtpG expression. Both strains supported robust overexpression of seHtpG without significant differences. High protein yield was achieved under both induction conditions: 3 hours at 37°C or overnight at 16°C following IPTG addition. Our findings indicate that cyanobacterial HtpG, as exemplified by seHtpG, is well-suited for efficient recombinant overexpression in *E. coli*.

**Figure 2.**
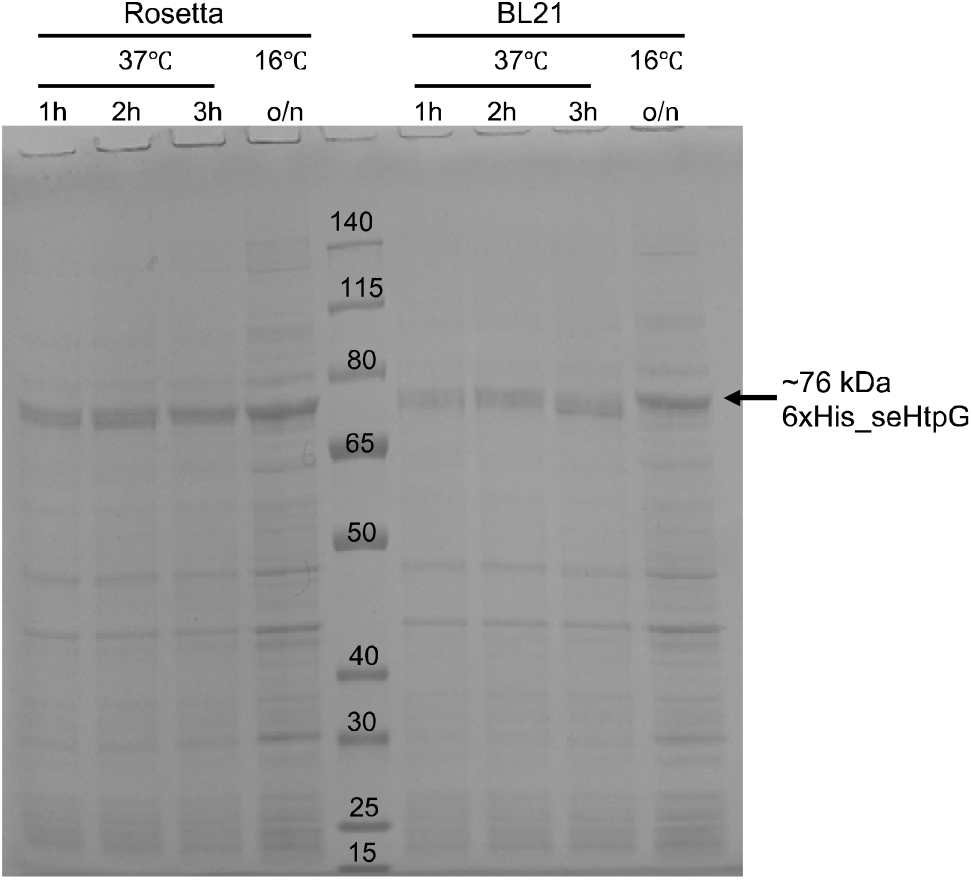
The recombinant overexpression level of seHtpG visualized by SDS-PAGE gel analysis with Coomassie staining. The protein makers (in kDa) were loaded into the middle lane.

### 3.3 Purification of recombinant seHtpG

The purification of recombinant seHtpG was conducted through a three-step process. First, the overexpressed protein was subjected to Ni-NTA affinity chromatography, followed by anion exchange chromatography and then size-exclusion chromatography. The N-terminal 6xHis tag was removed after affinity purification via cleavage with TEV protease. The elution profile from anion exchange chromatography is shown in Fig. 3A. The protein eluted as a single peak at a conductivity of 31 mS/cm, which corresponds to an approximate salt concentration of 300 mM KCl. This finding indicates that the surface of seHtpG is highly negatively charged, consistent with its low isoelectric point (pI ≈ 4.9). Furthermore, the protein’s propensity to bind to the anion exchange column is likely enhanced by its dimerization, as confirmed by subsequent size-exclusion chromatography and analytical ultracentrifugation (AUC).

**Figure 3.**
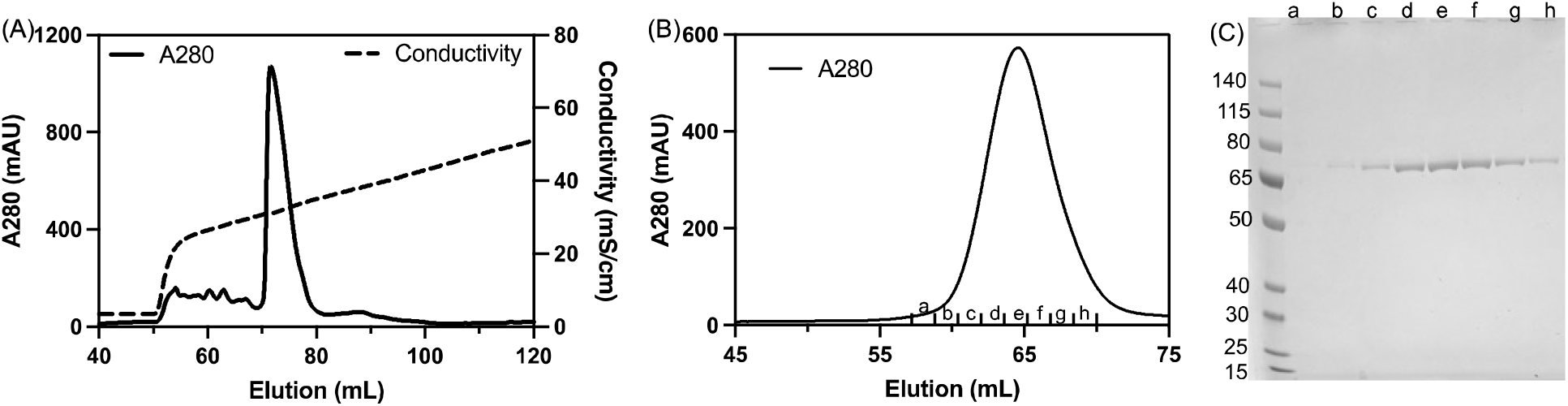
Purification of recombinant seHtpG (A) Elution profile of seHtpG from anion exchange chromatography. (B) Elution profile of seHtpG from size exclusion chromatography. (C) SDS-PAGE analysis of seHtpG fractions collected from size exclusion chromatography shown in (B). The first lane on the left contains the molecular weight marker (in kDa). Lanes a–h correspond to seHtpG sample fractions collected from size exclusion chromatography in (B).

To achieve high purity, the final step in the purification of seHtpG involved size-exclusion chromatography, which also served the purpose of buffer exchange and putting the protein into the storage buffer (40 mM Hepes pH 7.5, 150 mM KCl, 0.5 mM TCEP). The elution profile from size-exclusion chromatography is shown in Fig. 3B. Purity exceeding 99% was confirmed by SDS-PAGE, with no contaminating proteins visible (Fig. 3C). The observed molecular weight on SDS-PAGE fell between 65 kDa and 80 kDa, consistent with the theoretical molecular weight of 72.6 kDa for monomeric seHtpG. The size-exclusion chromatography peak eluted at 65 mL with a flow rate of 0.5 mL/min (Fig. 3B), corresponding to a molecular weight close to that of the 158 kDa IgG standard provided by the column manufacturer (GE Life Sciences). This gel-filtration result indicates that recombinant seHtpG exists predominantly as a dimer, consistent with other Hsp90 homologs. A high yield of ∼20 mg/L was achieved following the three-step purification process.

### 3.4 seHtpG forms a dimer

To assess the oligomeric state of purified seHtpG, we performed analytical ultracentrifugation (AUC) as described in the Methods section. The sedimentation velocity experiment confirmed that seHtpG is dimeric, with a sedimentation coefficient of sw=5.458□S, sw(20,w)=5.743S, corresponding to an approximate molecular weight of 130 kDa (Fig. 4). Other Hsp90 orthologs exhibit similar sedimentation coefficients, including 5.96 S for *LbHsp90* [18], 6.10 S for porcine brain Hsp90 [19], 6.0 S for human mitochondrial Trap1 [20], and 5.6 S for *E. coli* HtpG [21]. The AUC results for seHtpG being a dimer are consistent with findings from size-exclusion chromatography.

**Figure 4.**
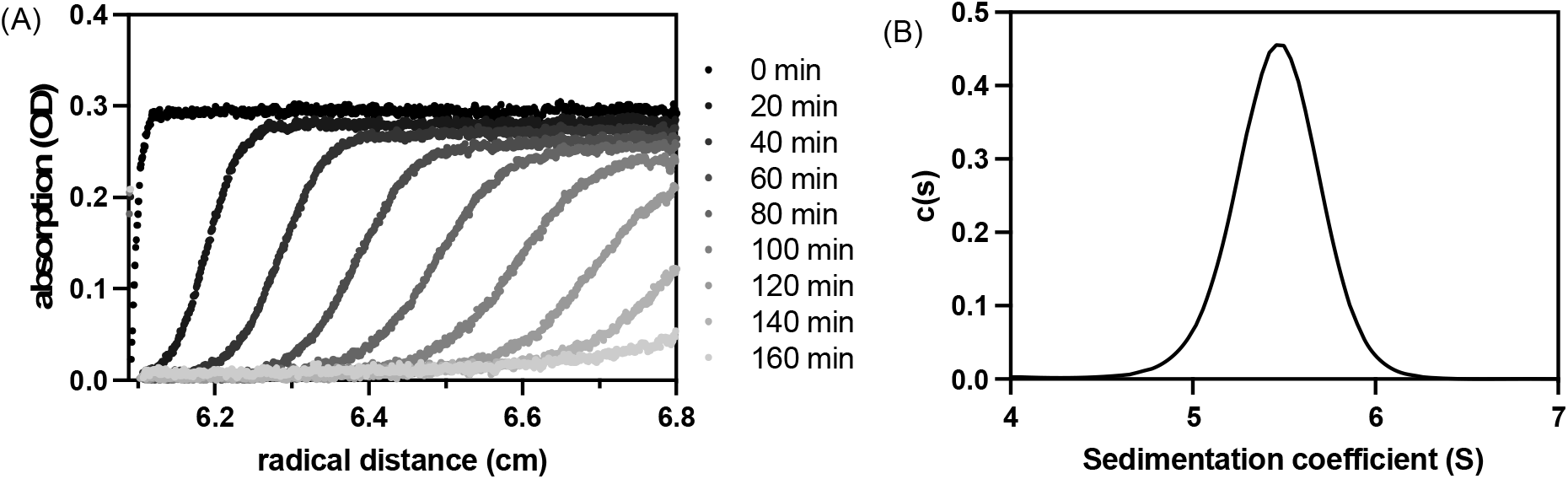
Analytical ultracentrifugation (AUC) analysis of seHtpG. (A) Representative sedimentation velocity traces for seHtpG depicted every 20 minutes for the sake of clarity. Absorbance at 280 nm was monitored for 4 µM seHtpG. (B) Sedimentation coefficient distribution for seHtpG, determined from a global fit of the sedimentation velocity traces.

### 3.5 ATPase activity of recombinant seHtpG

We confirmed that the purified recombinant seHtpG is active by measuring its ATPase activity. Members of the Hsp90 family are characterized by intrinsic ATPase activity [22]. To investigate the biochemical function of seHtpG, enzyme-coupled ATPase assays were conducted at various seHtpG concentrations in the presence of 1 mM ATP and 1 mM Mg^2+^. ATPase activity was assessed by monitoring the rate of NADH consumption, which is coupled to ADP release from ATP hydrolysis. The decrease in NADH absorbance at 340 nm (Fig. 5A) indicates that seHtpG is functional and actively hydrolyzes ATP. The hydrolysis rate was found to depend linearly on the concentration of seHtpG (Fig. 5B). To determine the average ATP hydrolysis rate, the ADP release rate was normalized to the seHtpG monomer concentration, then averaged across three different concentrations. The final normalized ATP hydrolysis rate was 1.81 ± 0.08 ATP/min at 37°C, which is higher than the reported ATPase activities of 0.4 ATP/min for Hsp82 and 0.48 ATP/min for *E. coli* HtpG [23], 0.8 ATP/min for *Mycobacterium tuberculosis* HtpG [24] at the same temperature.

**Figure 5.**
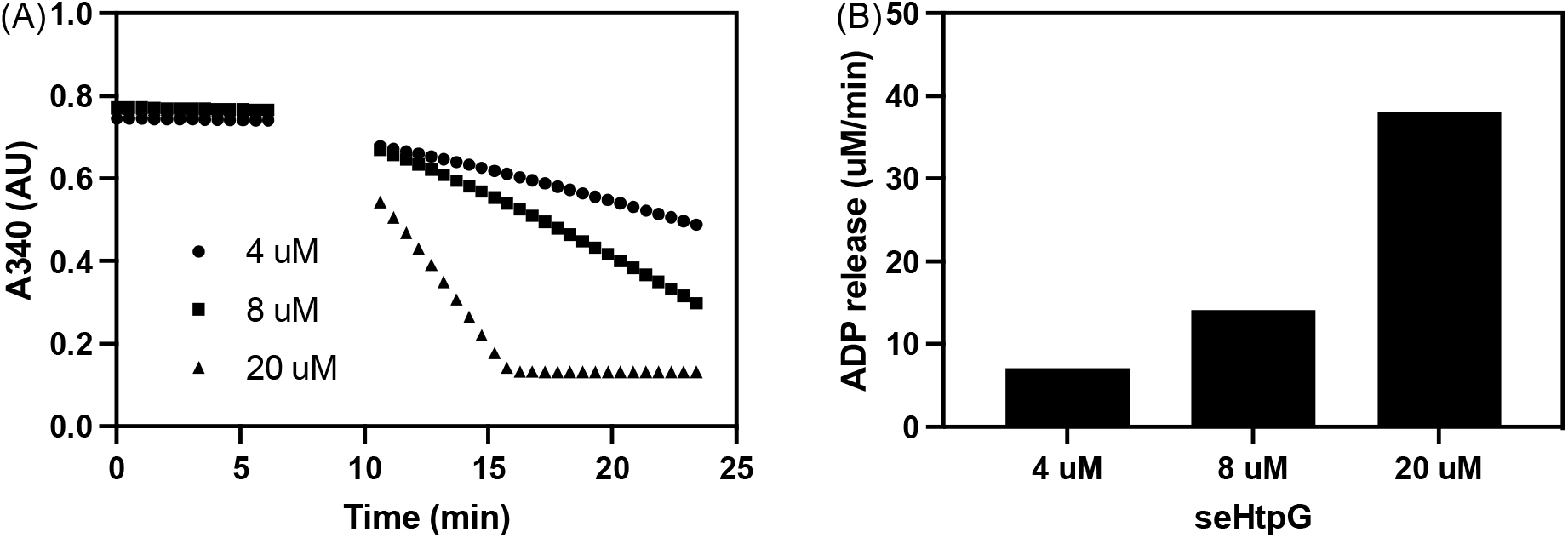
ATP hydrolysis activity of seHtpG measured by NADH based enzymatic coupled assay. (A) Protein concentration dependence of ADP release over time by monitoring NADH absorbance at 340 nm. The reaction mixtures were pre-equilibrated at 37°C for 7 minutes before ATP addition. (B) ADP release rates at different seHtpG concentrations, determined by fitting the linear regions of the curves in (A) after ATP addition. The final ATP hydrolysis rate for seHtpG was calculated as 1.81 ± 0.08 ATP/min by first normalizing the ADP release rate by protein concentration and then averaging across three protein concentrations.

## 4. Conclusion

The role of the molecular chaperone HtpG in cyanobacterial physiology, photosynthesis, and stress response remains poorly understood. Here, we demonstrate that HtpG from the cyanobacterium *Synechococcus elongatus* PCC 7942 (seHtpG) can serve as a model protein for the biochemical and biophysical investigation of cyanobacterial HtpG. While seHtpG exhibits conserved sequence and structural features typical of all Hsp90 homologs, it also presents unique characteristics specific to cyanobacteria, such as a sequence insertion in the middle domain. We established a protocol to recombinantly overexpress and purify seHtpG to apparent homogeneity with high yield, making it suitable for biochemical and high-resolution structural studies using X-ray crystallography, cryo-electron microscopy, and NMR. The purified seHtpG forms a dimer and is functionally active, hydrolyzing ATP at a rate of 1.8 ATP/min, faster than that of *E. coli* HtpG. Overall, our work lays a foundation for further investigation into the structure, function, and client interactions of cyanobacterial HtpG in *S. elongatus* and related cyanobacteria.

## Acknowledgements

The project is supported by start-up funding from the University of Maryland College Park and The Institute for Bioscience and Biotechnology Research Seed Grant funded by the University of Maryland Strategic Partnership: MPowering the State.

## CRediT authorship contribution statement

**Liqun Jiang:** Conceptualization, Data curation, Formal analysis, Investigation, Methodology, Project administration, Resources, Validation, Visualization, Writing – original draft, Writing – review and editing. **Ibrahim D. Boyenle:** Investigation, Methodology, Writing – review and editing. **Nicolas Delaeter:** Investigation, Methodology, Writing – review and editing. **Yanxin Liu:** Conceptualization, Funding acquisition, Investigation, Project administration, Supervision, Writing – original draft, Writing – review and editing.

## Declaration of generative AI and AI-assisted technologies in the writing process

During the preparation of this work the authors used ChatGPT in order to check the grammar and polish the sentences. After using this tool, the authors reviewed and edited the content as needed and take full responsibility for the content of the publication.

